# Invasive atypical non-typhoidal *Salmonella* serovars in The Gambia

**DOI:** 10.1101/2021.02.18.431831

**Authors:** Abdoulie Kanteh, Abdul Karim Sesay, Nabil-Fareed Alikhan, Usman Nurudeen Ikumapayi, Rasheed Salaudeen, Jarra Manneh, Yekini Olatunji, Andrew J Page, Grant Mackenzie

**Affiliations:** Medical Research Council Unit The Gambia at the London School of Hygiene and Tropical Medicine, Atlantic Boulevard, Fajara, PO Box 273, Banjul, The Gambia; Faculty of Infectious and Tropical Diseases, London School of Hygiene and Tropical Medicine, Keppel Street, London, WC1E 7HT, United Kingdom; Murdoch Children’s Research Institute, Royal Children’s Hospital Flemington Road, Parkville Victoria 3052 Australia; Department of Paediatrics, University of Melbourne, Melbourne, Australia; Quadram Institute Bioscience, Norwich Research Park, Norwich, Norfolk, NR4 7UA, UK

**Keywords:** Invasive non-typhoidal salmonella, Whole genome sequencing, Cytolethal distending toxin gene, atypical serovar

## Abstract

**Background:** Invasive non-typhoidal *Salmonella* (iNTS) disease continues to be a significant public health problem in sub-Saharan Africa. Common clinical misdiagnosis, antimicrobial resistance, high case fatality and lack of a vaccine make iNTS a priority for global health research. Using whole genome sequence analysis of 164 invasive *Salmonella* isolates obtained through population-based surveillance between 2008 and 2016, we conducted genomic analysis of the serovars causing invasive *Salmonella* diseases in rural Gambia.

**Results:** The incidence of iNTS varied over time. The proportion of atypical serovars causing disease increased over time from 40% to 65% compared to the typical serovars Enteritidis and Typhimurium decreasing from 30% to 12%. Overall iNTS case fatality was 10% with 10% fatality in cases of atypical iNTS. Genetic virulence factors were identified in 14/70 (20%) typical serovars and 45/68 (66%) of the atypical serovars and were associated with: invasion, proliferation and/or translocation (Clade A); and host colonization and immune modulation (Clade G). Among Enteritidis isolates, 33/40 were resistant to ≥4 the antimicrobials tested, except for ciprofloxacin, to which all isolates were susceptible. Resistance was low in Typhimurium isolates, however, all16 isolates were resistant to gentamicin.

**Conclusion:** The increase in incidence and proportion of iNTS disease caused by atypical serovars is concerning. The increased proportion of atypical serovars and the high associated case fatality may be related to acquisition of specific genetic virulence factors. These factors may provide a selective advantage to the atypical serovars. Investigations should be conducted elsewhere in Africa to identify potential changes in the distribution iNTS serovars and the extent of these virulence elements.

## Introduction

The species *Salmonella enterica* (*S. enterica*) is a phenotypically diverse Gram-negative bacterial species, consisting of more than 2,600 serovars. Some serovars are implicated in life-threatening systemic infections and are host-restricted to humans^1^. These include *Salmonella enterica* serovar Typhi and *Salmonella enterica* serovar Paratyphi (*S. Paratyphi* A-C). In contrast, non-typhoidal *Salmonella* species infect both humans and animals^2^; *Salmonella enterica* serovar Typhimurium and *Salmonella enterica* serovar Enteritidis are the most commonly reported in association with *Salmonella* gastroenteritis^3^. Globally, these serovars are responsible for circa 75 million cases and 27,000 deaths annually^3^.

In sub-Saharan Africa, in addition to causing gastroenteritis, non-typhoidal *Salmonella* (NTS) cause life-threatening infections including septicaemia, pneumonia and meningitis^4^. Circa 3.4 million cases of invasive *Salmonella* caused by NTS (iNTS) are reported annually, with Typhimurium and Enteritidis being responsible for 80 - 90% of these cases^5^. The majority of these infections affect children, and are often associated with Human Immunodeficiency Virus (HIV) infection, prior malarial infection, severe anaemia or malnutrition, and case fatality of up to 25%^6–9^. In adults, HIV infection is associated with iNTS disease and case fatality up to 50% has been reported^7–9^. In some parts of Africa, the burden of iNTS disease is higher than that of pneumococcus, infecting tens of thousands of people^7–9^. In The Gambia, iNTS disease in children ranks third after *Streptococcus pneumoniae* and *Staphylococcus aureus* as a cause of invasive bacterial disease^10^. Despite the burden of this disease in our setting, the genomic epidemiology of NTS is still poorly understood.

Susceptibility to invasive *Salmonella* disease could be attributed to host genetic background and immunological status^4^. However, some serovars are known to cause bacteraemia more frequently than others, signifying the importance of pathogen characteristics. For example, a high burden of invasive disease caused by a specific genotype of *S.* Typhimurum has been associated with host adaptation as a result of extensive genomic degradation and acquisition of resistance genes^11^. In addition, the virulence factor cytolethal distending toxin gene *(CdtB)* is known to contribute to variation in disease severity in some NTS serovars^12^. The *CdtB* gene, which was thought to be unique to *Salmonella* Typhi, has been associated with increased host colonization, tumorigenesis, neoplastic lesions^13^ and DNA damage similar to that caused by serovar Typhi^13^. The presence of the gene in Typhi is associated with host immune modulation as well as persistence of the pathogen in *vivo*^12^. Recently, the presence of *CdtB* has also been documented in NTS serovars and is believed to be clade associated^12^. Thus, the presence of this virulence gene in NTS serovars could influence the virulence of these strains.

During population-based invasive bacterial disease surveillance in rural Gambia between 2008 and 2016, we observed changes in the incidence, case fatality, and distribution of iNTS serovars. Surveillance in the same location from 2000 to 2004 documented Enteritidis and Typhimurium as the dominant iNTS serovars^14^. Although shifts in *Salmonella* serovar prevalence and dominance have been documented in The Gambia and elsewhere in the world^14,15,16^, the genomic characteristics and epidemiological factors responsible for this shift are unclear. We used whole genome sequencing and bioinformatic analyses to investigate changes in pathogen characteristics between 2008 and 2016.

## Material and methods

### Disease surveillance

The surveillance methodology has been previously described^17^. We conducted population-based surveillance for invasive bacterial disease in individuals aged 2 months and older resident in the Basse Health and Demographic Surveillance System in Upper River Region, The Gambia^17^. We used standardised criteria to identify and investigate patients presenting with suspected pneumonia, septicaemia, or meningitis to all health facilities in the study area between May 12, 2008 and December 31, 2016. Blood, cerebrospinal fluid (CSF), and lung aspirates (LA) were collected according to standardised criteria and we used conventional microbiological methods to culture and identify bacterial pathogens. Gram negative isolates were identified as *Salmonella* biochemically using a commercial kit (Analytic Profile Index 20E) and antimicrobial susceptibility testing was done using the disk diffusion method and following CLSI reference thresholds^18^.

### Domestic animal ownership

Given that NTS also infects domestic animals, they can represent an important route of transmission. Data from the Global Enteric Multicentre Study^19^ collected in the study area between 2007 and 2012 were used to compare changes in the prevalence of domestic animal ownership and invasive *Salmonella* over time..

### Sample population

We analysed 164 *Salmonella* genomes from isolates obtained from blood, CSF or LA samples collected during the surveillance. We extracted genomic DNA from the isolates that was sent to the Wellcome Sanger Institute, United Kingdom for whole genome sequencing.

### Quality Control, Assembly and Resistance genes

Extracted DNA was sequenced using the lllumina Hiseq 2500 platform, to produce sequencing reads of 125 base pairs in FASTQ format ^20^, with a minimum target depth coverage of 50X. The reads and genomes were quality checked using FASTQC (v0.11.5) and an in-house pipeline, with manual review. The reads were of high quality with an average Phred score of 30 and thus did not require any trimming. Spades (v3.13.1) was used to perform *de novo* assembly with default settings^21^ to produce draft assemblies in FASTA format. Quast (v5.0.2)^22^ was used to assess the quality of assemblies. Contigs shorter than 300bp were removed from the assemblies as per Page *et al*.,^23^. Four genomes were significantly larger (six Mbases) than the rest of the genomes indicating contamination and were therefore removed from the analysis.

We used Abricate (v0.9.8) to identify antimicrobial resistance genes, plasmids and virulence genes for each assembly using the comprehensive antimicrobial resistance database (CARD)^24^ (downloaded 24-10-2019), Resfinder^25^ (downloaded 10-9-2019), PlasmidFinder^26^ (downloaded 10-9-2019) and the virulence factor database (VFDB)^27^ (downloaded 18-09-2019). A minimum nucleotide identity and coverage of 98% was used for all databases. Virulence factors universally present in *Salmonella* were excluded. The multilocus sequence type (MLST) of each draft genome was predicted using mlst (v2.8) with default settings against the *Salmonella enterica* MLST scheme in the PubMLST database^28^.

### Phylogenetic analysis

Sequencing reads were mapped to the *Salmonella enterica* serovar Typhimurium LT2 reference genome (accession number GCF_000006945.2) using Snippy (v4.0.7) with default settings. Single nucleotide polymorphisms (SNPs) from the core genome alignment were used to construct a maximum likelihood phylogenetic tree using the general time-reversible model with IQTREE (v1.3.11.1)^29^ and 1000 bootstrap for branch length. Interactive Tree of Life (ITOL) (v5)^30^ was used to visualise and annotate the phylogenetic tree. Where particular serovars appeared to have developed into an outbreak they were analysed phylogenetically with other isolates from outside our study. In addition, when genotypes (or STs) were identified that were known to be resitricted elsewhere in the world, phylogenetic comparisons were made to determine whether they were related.

### Pan and accessory genome analysis

We used Prokka (v1.13.3)^31^ to annotate and predict coding genes from the assembled genomes using *S.* Typhimurium LT2 protein sequences from GenBank to provide high quality species-specific gene name annotation. The resulting GFF3 files were used as input to Roary (v3.13.2)^32^ to generate a pan-genome, producing an analysis of the core and accessory genome.

### Statistical analysis

Summary statistics were prepared using proportions for categorical and mean/median/range for continuous variables including demographic and baseline characteristics. We used Fisher’s exact test for associations between categorical variables. All data management and statistical analyses were performed using the R statistical package.

## Results

### Demographic data

Between 2008 and 2016, 22,305 patients were enrolled in the surveillance with 20,199 microbiological cultures, an average 2,244 per year (range: 1,047 – 2,370) (Table 1). Patient characteristics are shown in Table 2. From all cultures collected, 164 *Salmonella* isolates were obtained from 157 patients. Patient age ranged from 3 days to 42 years with children aged <5 years representing more than 90% (n=145) of the cases. By sample type, 157 isolates were from blood, six from CSF and one from LA. Six patients had isolates detected from more than one clinical sample type.

**Table 1.** Numbers of patients enrolled, blood cultures collected, and *Salmonella* isolates detected each year

| Year | Total enrolled | Total blood cultures taken | Enteritidis | Typhimurium | Typhi | Atypical | Paratyphi | Total |
| --- | --- | --- | --- | --- | --- | --- | --- | --- |
| 2008 | 1212 | 1047 | 0 | 0 | 3 | 3 | 1 | 7 |
| 2009 | 2099 | 1898 | 1 | 2 | 1 | 4 | 0 | 8 |
| 2010 | 1869 | 1605 | 6 | 3 | 1 | 2 | 0 | 12 |
| 2011 | 2688 | 2385 | 23 | 2 | 4 | 4 | 0 | 33 |
| 2012 | 2899 | 2592 | 7 | 3 | 7 | 20 | 0 | 37 |
| 2013 | 2580 | 2200 | 2 | 3 | 5 | 17 | 0 | 27 |
| 2014 | 2707 | 2536 | 7 | 3 | 3 | 9 | 0 | 22 |
| 2015 | 3742 | 3566 | 3 | 2 | 2 | 5 | 0 | 12 |
| 2016 | 2509 | 2370 | 0 | 1 | 1 | 4 | 0 | 6 |
| Total | 22305 | 20199 | 49 | 19 | 27 | 68 | 1 | 164 |

**Table 2.** Summary of baseline patient characteristics.

| Variable | Characteristic | N (%) |
| --- | --- | --- |
| Sex | Male | 84 (53.5) |
|  | Female | 73 (46.5) |
| Diagnosis | Pneumonia | 93 (59.2) |
|  | Meningitis | 11 (7.0) |
|  | Septicaemia | 46 (29.3) |
|  | Other focal sepsis | 6 (3.8) |
|  | Other | 1 (0.6) |
| Disease Outcome | Dead | 16 (10.2) |
|  | Discharged and/or recovered | 111 (70.7) |
|  | Not admitted | 22 (14.0) |
|  | Absconded | 1 (0.6) |
|  | Transferred | 6 (3.8) |
|  | Missing | 1 (0.6) |
| Age range | 0-5 yrs | 144 (91.7) |
|  | 6-15 yrs | 7 (4.5) |
|  | >15 yrs | 6 (3.8) |
| Nutritional status | Acute malnutrition | 51 (32.5) |
|  | Moderate acute malnutrition | 32 (20.4) |
|  | Well nourished | 64 (40.8) |
|  | Missing | 10 (6.4) |
| Reside with the surveillance area | Yes | 136(86.6) |
|  | No | 21(13.4) |
| Sample type | Blood | 157 (95.7) |
|  | Cerebospinal fluid | 6 (3.7) |
|  | Lung Aspirate | 1 (0.6) |
| Infection rate by serotype | Enteritidis | 47 (29.9) |
|  | Typhimurium | 18 (11.5) |
|  | Typhi | 27 (17.2) |
|  | Paratyphi C | 1 (0.6) |
|  | Atypical | 64 (40.8) |

**Table 3.** Summary of resistance and plasmid genes in each serovar.

| Clade | Serovar | Gene Name | Total (%) | Plasmid genes | Total (%) |
| --- | --- | --- | --- | --- | --- |
| A | Dublin | fosA7_1 | 1/14 (7.1) | IncFII(S)_1 | 14/14 (100) |
|  |  |  |  | IncI1_1_Alpha | 1/14 (7.1) |
|  |  |  |  | IncX1_1 | 14/14 (100) |
| B | Enteritidis | aph(3'')-Ib_5 | 45/49 (91.8) | ColpVC_1 | 1/49 (2.1) |
|  |  | aph(6)-Id_1 | 45/49 (91.8) | IncFIB(S)_1 | 2/49 (4.1) |
|  |  | blaTEM-1B_1 | 49/49 (100) | IncFII(S)_1 | 2/49 (4.1) |
|  |  | catA1_1 | 46/49 (93.8) | IncI1_1_Alpha | 47/49 (95.9) |
|  |  | dfrA7_5 | 46/49 (93.8) | IncQ1_1 | 45/49 (91.8) |
|  |  | sul1_5 | 46/49 (93.8) | rep21_9_rep(pKH12) | 2/49 (4.1) |
|  |  | sul2_6 | 45/49 (91.8) |  |  |
|  |  | tet(B)_2 | 46/49 (93.8) |  |  |
| C | Typhimurium | aph(3'')-Ib_5 | 3/19 (15.8) | IncFIB(S)_1 | 18/19 (94.7) |
|  |  | aph(6)-Id_1 | 3/19 (15.8) | IncFII(S)_1 | 18/19 (94.7) |
|  |  | blaTEM-1B_1 | 3/19 (15.8) | IncQ1_1 | 1/19 (5.3) |
|  |  | catA1_1 | 3/19 (15.8) |  |  |
|  |  | dfrA7_5 | 3/19 (15.8) |  |  |
|  |  | sul1_5 | 3/19 (15.8) |  |  |
|  |  | sul1_3 | 3/19 (15.8) |  |  |
|  |  | fosA7_1 | 1/19 (5.3) |  |  |
|  | Stanleyville | fosA7_1 | 2/2 (100) |  |  |
| D | Hofit | fosA7_1 | 1/1 (100) | IncFIB(S)_1 | 1/1 (100) |
|  |  |  |  | IncFII(S)_1 | 1/1 (100) |
| E | Virchow | no gene |  | pSL483_1 | 1/7 (14.3) |
| F | Typhi | catA1_1 | 2/27 (7.4) | no plasmid |  |
|  |  | dfrA7_5 | 2/27 (7.4) |  |  |
|  |  | sul1_5 | 2/27 (7.4) |  |  |
| G | Others: |  |  |  |  |
|  | Mountpleasant | fosA7_1 | 1/41 (100) | IncFII(S)_1 | 2/41 (4.8) |
|  | Senftenberg | fosA7_1 | 1/41 (100) | IncFII(pCoo)_1_pCoo | 1/41 (2.4) |
|  | Grumpensis | fosA7_1 | 1/41 (100) |  |  |
|  | Paratyphi C | fosA7_1 | 1/1 (100) | IncFIB(S)_1 | 1/1 (100) |
|  |  |  |  | IncFII(S)_1 | 1/1 (100) |

### Genomic analysis

MLST analysis revealed 31 distinct serovars and 45 sequence types (ST). We detected 27 serovars that were not Enteritidis, Typhimurium, Typhi or Paratyphi. We grouped these isolates and called them atypical serovars. A considerable proportion, 41% (n=68) of isolates were atypical. The atypical serovars most commonly isolated were Dublin (n=14) Virchow (n=7) and Poona (n=5). Enteritidis, Typhimurium and Typhi constituted 30% (n=49), 12% (n=19) and 16% (n=27) of the isolates, respectively. Only one isolate was S*almonella enterica* serovar Paratyphi C of ST 3039 (Figure 1).

**Figure 1.**
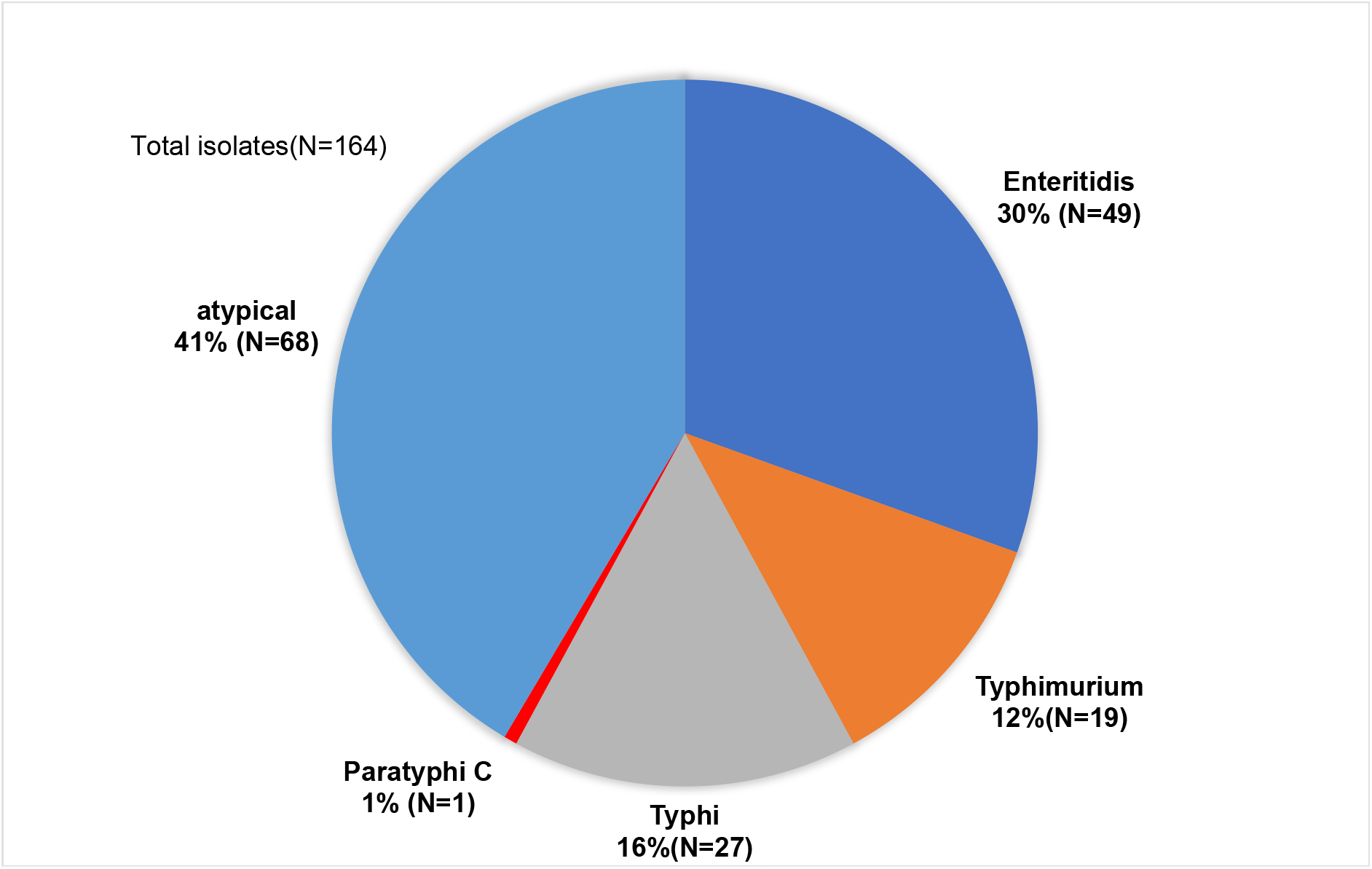
Breakdown of invasive *Salmonella* serovars isolated between 2008 and 2016 from patients in rural Gambia.

Of all the STs, ST11 was dominant, representing 30% (n=49) of the isolates, followed by ST2 which accounted for 16% (n=27). ST10 and ST19 represented 9% (n=14) and 8% (n=13) of the isolates respectively. Other STs included ST313 (n=4), ST3031 (n=3) and ST359 (n=3). Isolates of Typhimurium were represented by four STs: ST19, ST313, ST2988 and ST165. Serovars Virchow and Poona were represented by three and four STs, respectively. Some atypical serovars, including Bredeney, Give, Miami, Oranienburg, Overschie, Poona, Stanleyville and Virchow, were represented by two or more STs each. In contrast, serovars Enteritidis, Typhi and Dublin were represented by only one ST each: ST11, ST2 and S10 respectively (Figure 2).

**Figure 2.**
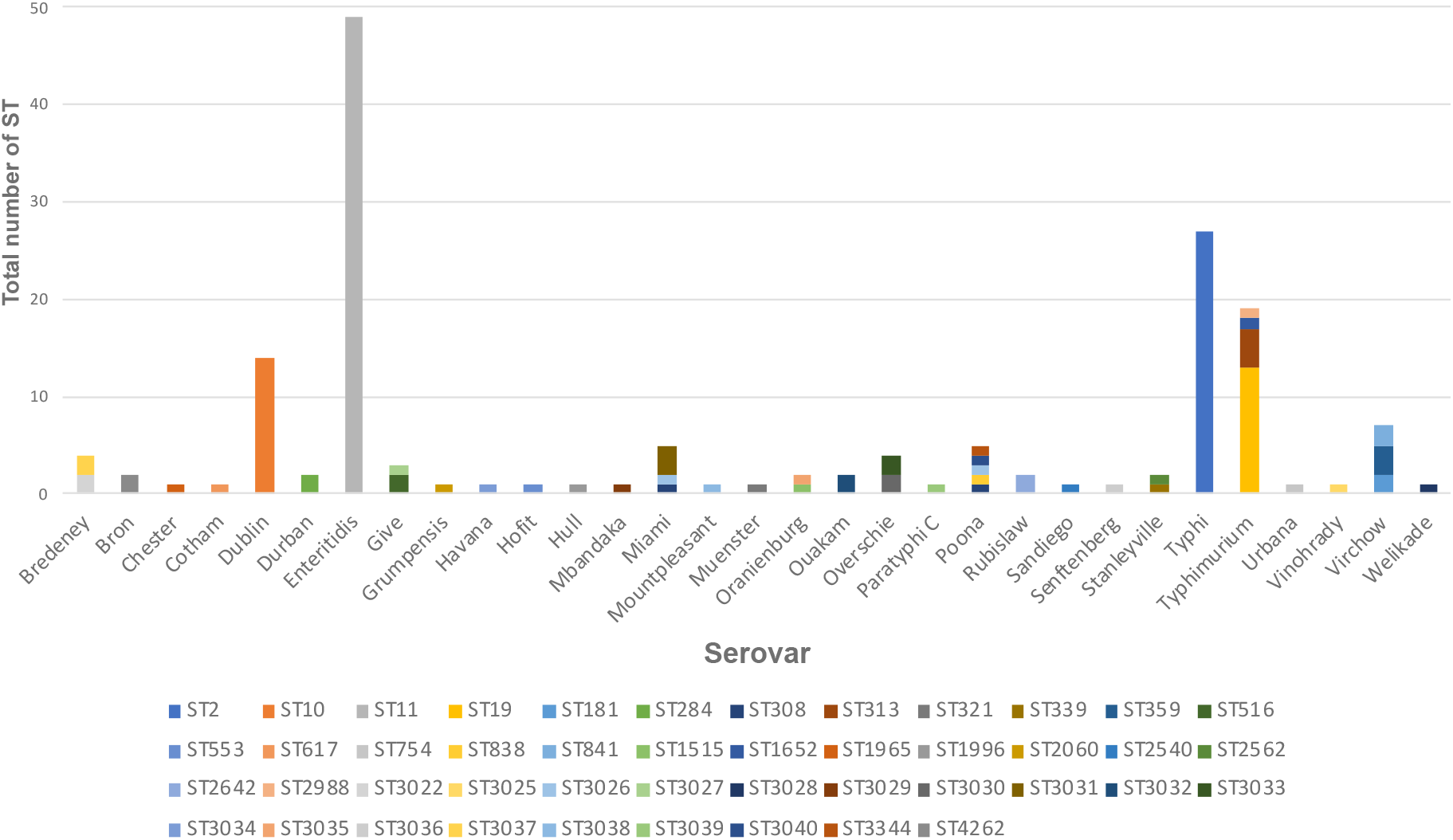
Representation of STs amongst invasive *Salmonella* serovars isolated between 2008 and 2016 from patients in rural Gambia.

### Distribution of *Salmonella* serovars over time

During 2000 to 2004 serovars Enteritidis (81%) and Typhimurium (8%) were the dominant iNTS serovars ^14^. Over the study period, we observed an increase in the proportion of atypical serovars (Figure 3). In 2008 and 2009, invasive *Salmonella* infection caused by atypical serovars accounted for the majority of cases compared with infection caused by Enteritidis and Typhimurium. However, this trend changed in 2011 when Enteritidis became predominant and accounted for about 80% of all *Salmonella* cases. A high proportion of atypical serovars was then observed between 2012 and 2014. Overall, from 2012 to 2014, atypical serovars were responsible for almost 50% of *Salmonella* infections. The major serovars within this group included Dublin, Bredeney, Miami and Overchie. From 2015 to 2016, we observed a further decline in the proportion of Enteritidis and Typhimurium serovars in the population, while atypical serovars were associated with over 50% of cases.

**Figure 3.**
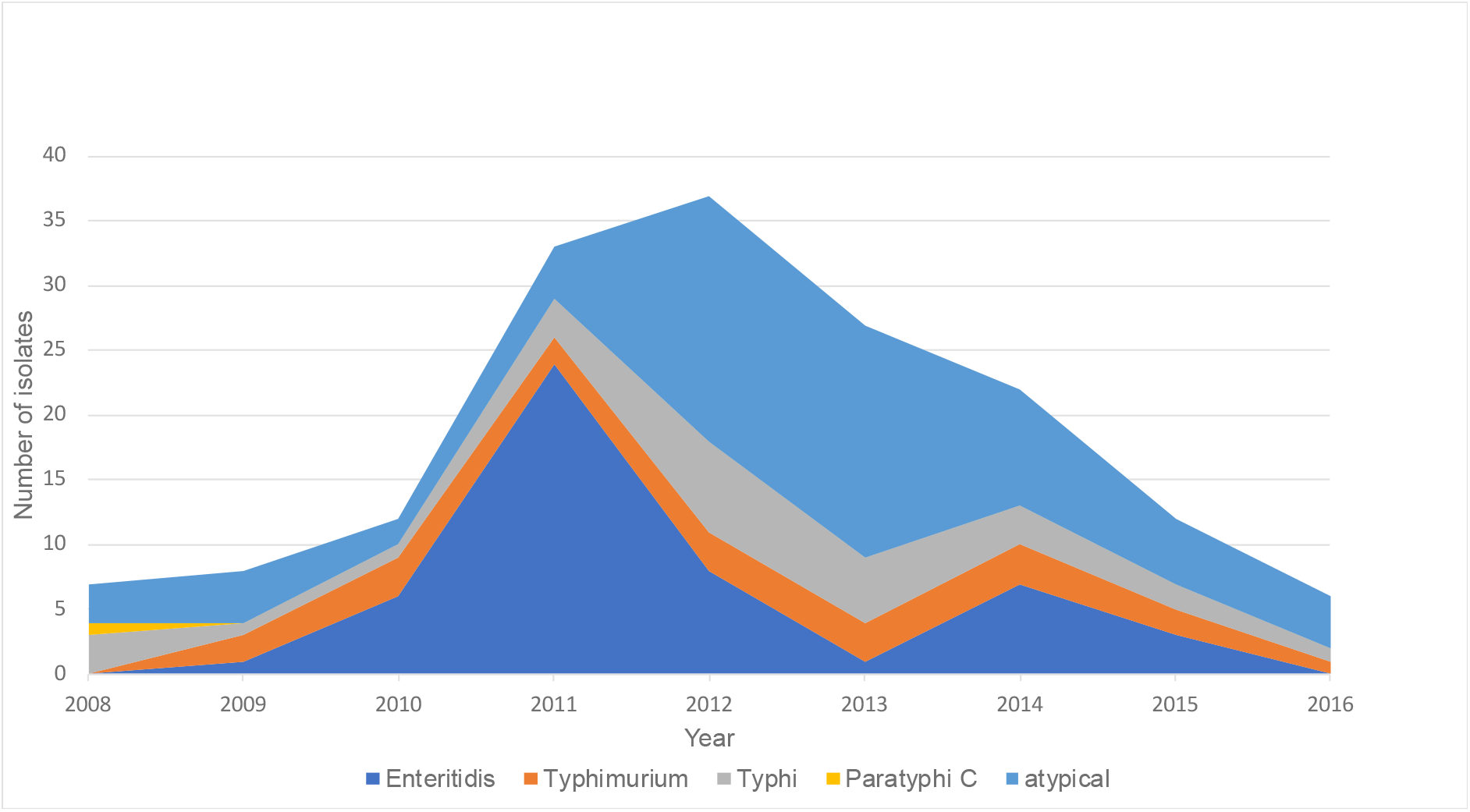
Case counts of each type of invasive *Salmonella* serovar in Basse, rural Gambia between 2008 and 2016.

### Incidence and case fatality rate

Amongst all cases of invasive *Salmonella* disease, case fatality rate was 10% (16/157). Case fatality for atypical serovars was 10% (7/68) and 12% (6/49) for Enteritidis. Typhi, Typhimurium and Paratyphic C were associated with only one death each. Amongst hospitalised patients, Enteritidis and atypical serovars accounted for 42% (32/77) and 31% (24/77) of cases while Typhi and Typimurium accounted for 16% (12/77) and 13% (10/77) of cases, respectively. Amongst atypical serovars, those with the cytolethal toxin gene *CdtB* were responsible for 10% (3/31) of all deaths while atypical seovars without the toxin gene accounted for 11% (4/37) of all deaths.

The majority of the patients (59%) had suspected pneumonia or septicaemia (29%). Of the 46 patients with septicaemia, 26 (56%) were infected with atypical serovars; Dublin, Overchie, Bredeney and Poona accounted for most of these cases. Overall, we did not find a statistically significant association between malnutrition and any specific serovar though this should be interpreted with caution due to small numbers. However, comparing typical vs atypical serovars, the proportion of children with severe acute malnutrition 19/32 (59%) appeared to be higher in the atypical group compared to Enteriditis 6/32 (18%), Typhimurium 3/32 (9%) or Typhi 4/32 (12%),p-value=0.05.

### Domestic animal ownership and prevalence of NTS over time

The prevalence of invasive *Salmonella* increased from 2007 to 2010 while domestic animal ownership by households remained constant throughout this period (Figure 4).

**Figure 4.**
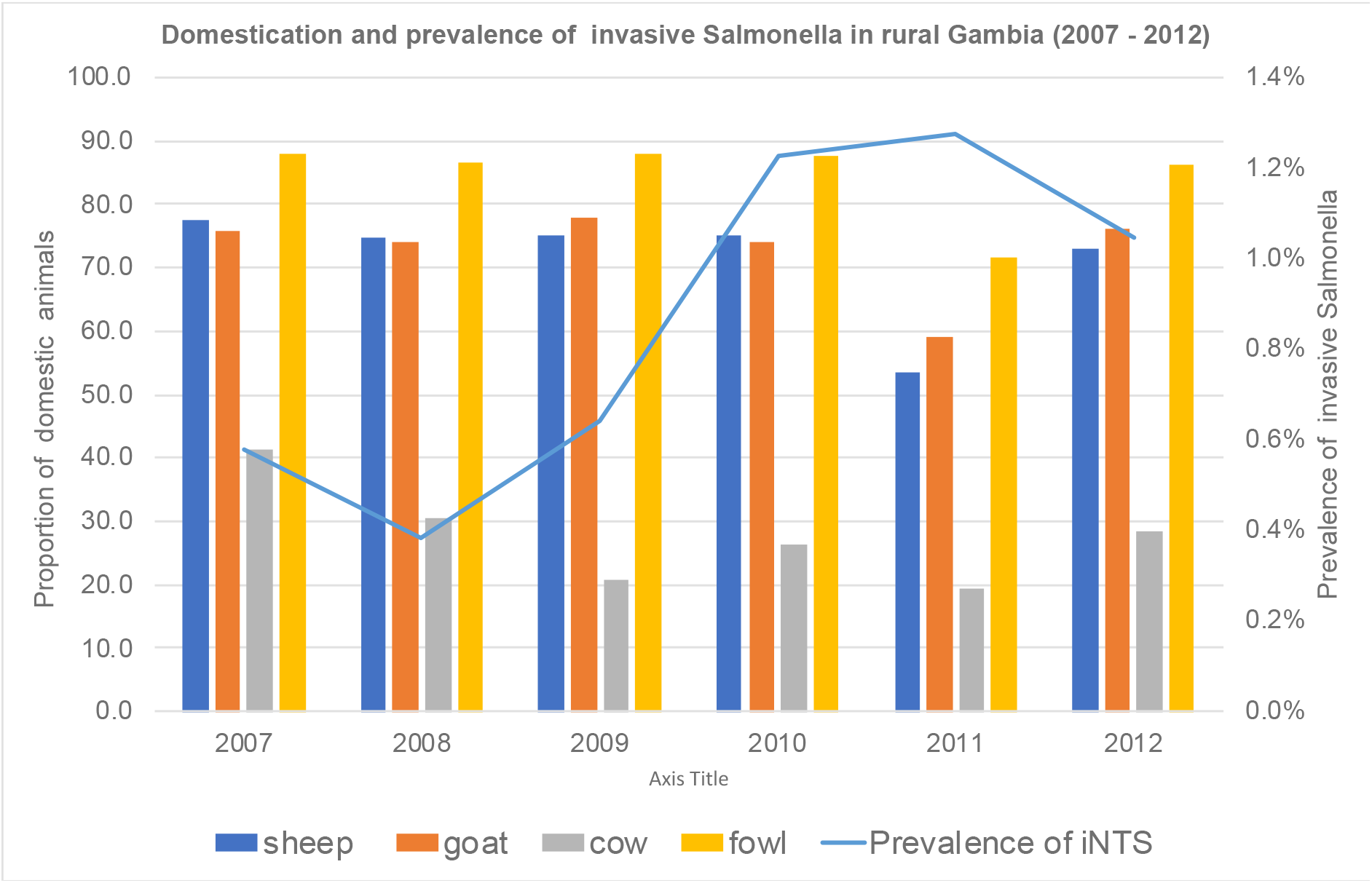
Relationship between invasive Salmonella disease incidence (blue line) and the proportion of different species of domestic animals reared in rural Gambia between 2007 and 2012.

### Phylogenetic analysis

We constructed a pan-*Salmonella* phylogenetic tree using single nucleotide polymorphisms (SNPs) generated from 3,331 sites in the core genome, excluding repeated regions and transposable elements. The tree resolved seven distinct clades. We named these clades A-G. Clade A and B were comprised of Dublin and Enteritidis serovars, respectively. Typhimurium clustered with Hull and Stanleyville in clade C. Clade D included serovars Hofit and Rubislaw while clade E was comprised only of Virchow isolates. All the Typhi isolates formed a distinct clade (clade F) and the remaining serovars formed a separate clade, clade G (Figure 5).

**Figure 5.**
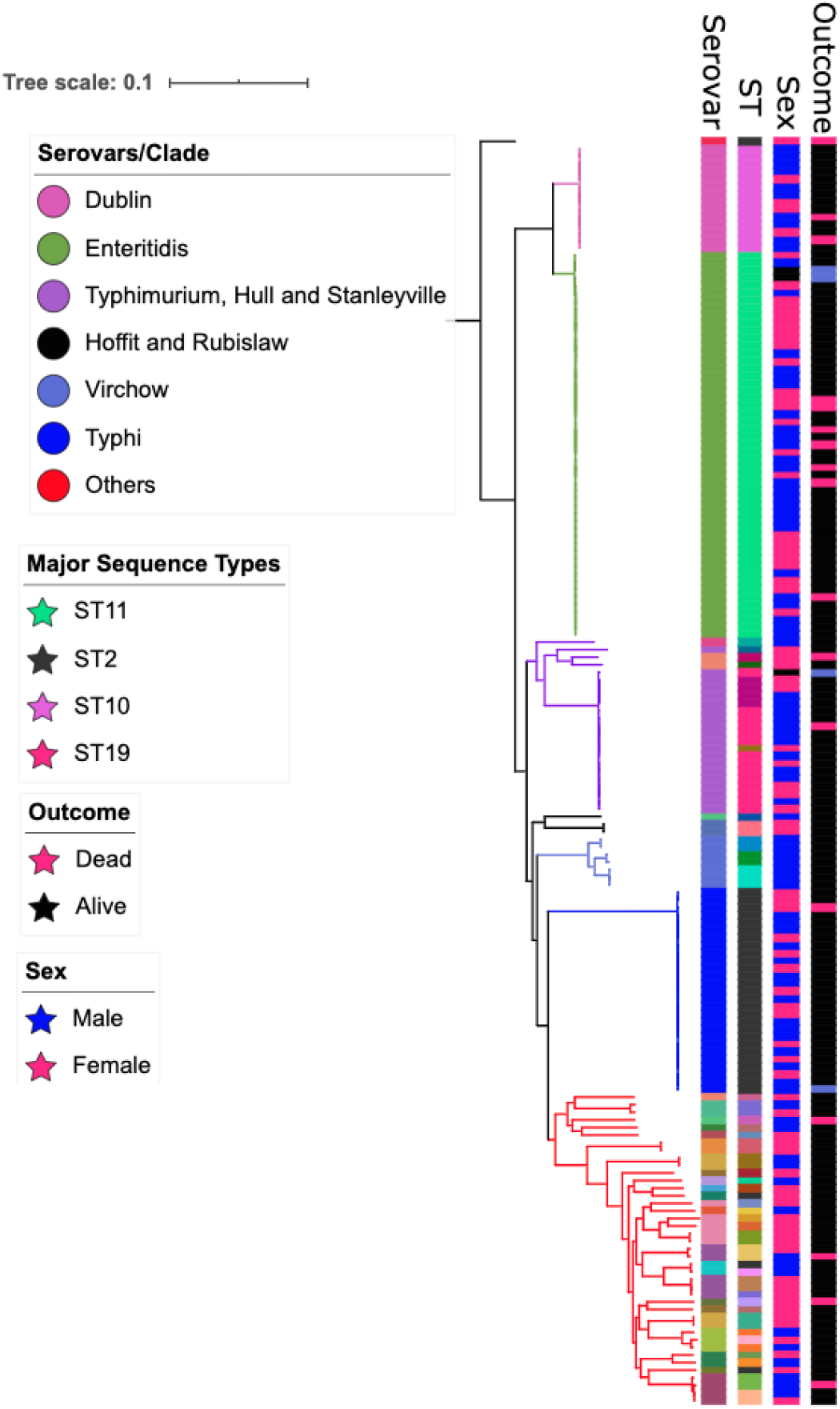
Maximum likelihood phylogenetic tree of 164 *Salmonella* genomes isolated from patients in rural Gambia between 2008 and 2016. Seven distinct clades were resolved from the tree and denoted by different colours (see legend). Metadata is shown alongside the phylogenetic and includes host sex and disease status. The serovars and most prevalent sequence types are annotated on the tree and denoted using different colours. The tree was rooted on the *Salmonella* Paratyphi C isolate.

### Genomic analysis of Enteritidis isolates

To understand the reason for the high proportion of Enteritidis between 2010 and 2011 we used phylogenetic analysis to compare the 2010 and 2011 Enteritidis genomes in our dataset with Enteritidis genomes collected in The Gambia before and after 2010. This analysis indicated a potential outbreak (Figure 6) with more than 70% (21/29) of the Enteritidis isolates collected during the surveillance in 2010 and 2011 clustered closely on the tree with short branch lengths, suggesting closely related strains circulating during this time frame.

**Figure 6.**
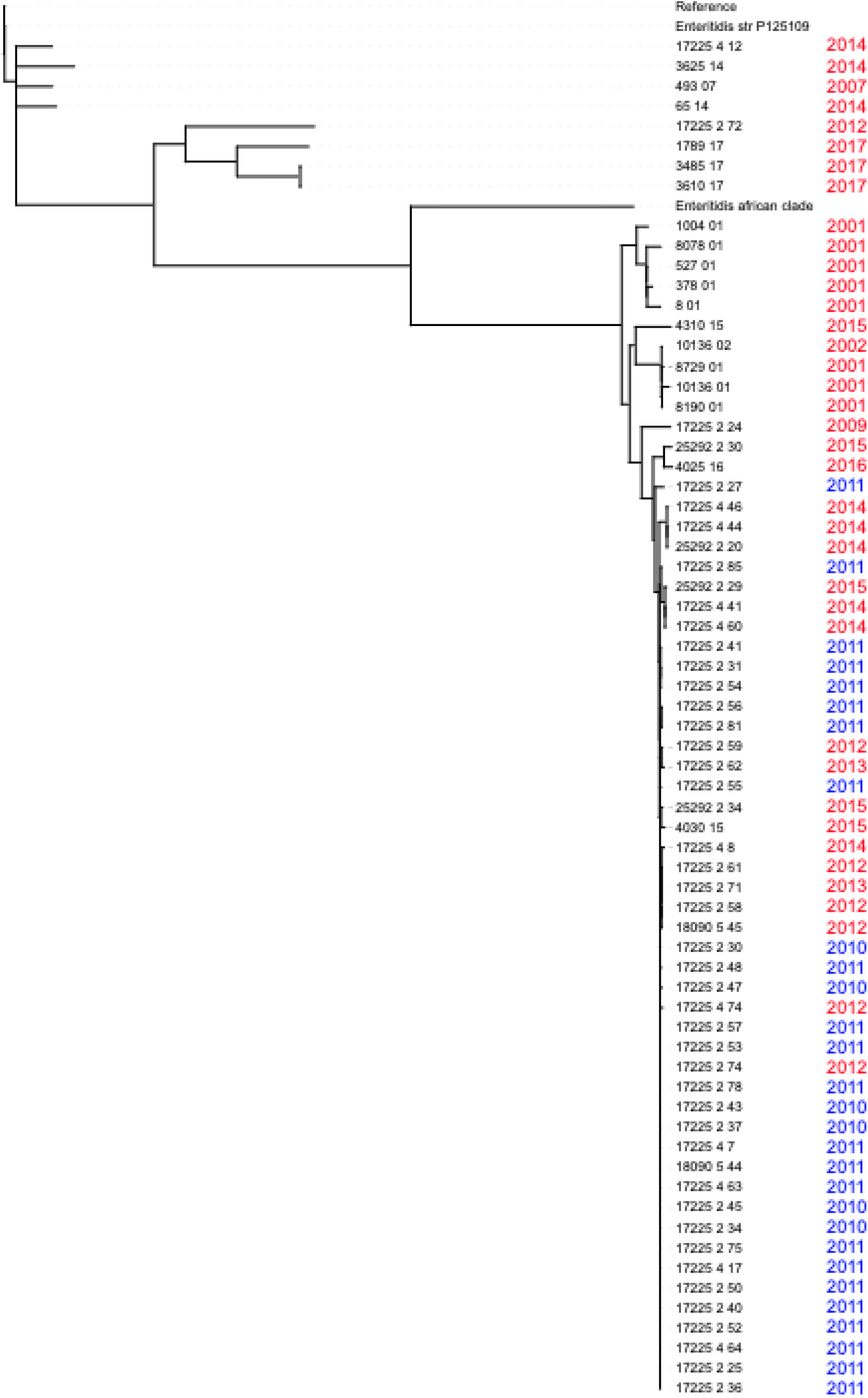
Phylogenetic tree of 49 *Salmonella* Enteritidis isolates collected during the surveillance period and 16 other isolates collected from The Gambia (both within the surveillance area and outside) at different time points. Isolates collected in the present study between 2010 and 2011 are colored blue and those collected before or after the surveillance period are coloured red. The tree is rooted on the *Salmonella* Typhimurium LT2 reference genome.

### Genomic analysis of S Typhimurium ST313 isolates

We found that five isolates had the ST313 genotype, which has been implicated as the causative agent of invasive *Salmonella* disease in Kenya and Malawi. For this reason, we used phylogenetic analysis to compare the ST313 isolates in our study with other global strains in Enterobase^33^. We found that the isolates circulating in The Gambia are of the lineage 1 type and different from the type circulating in Kenya and Malawi which are of the lineage 2 (Figure 7).

**Figure 7.**
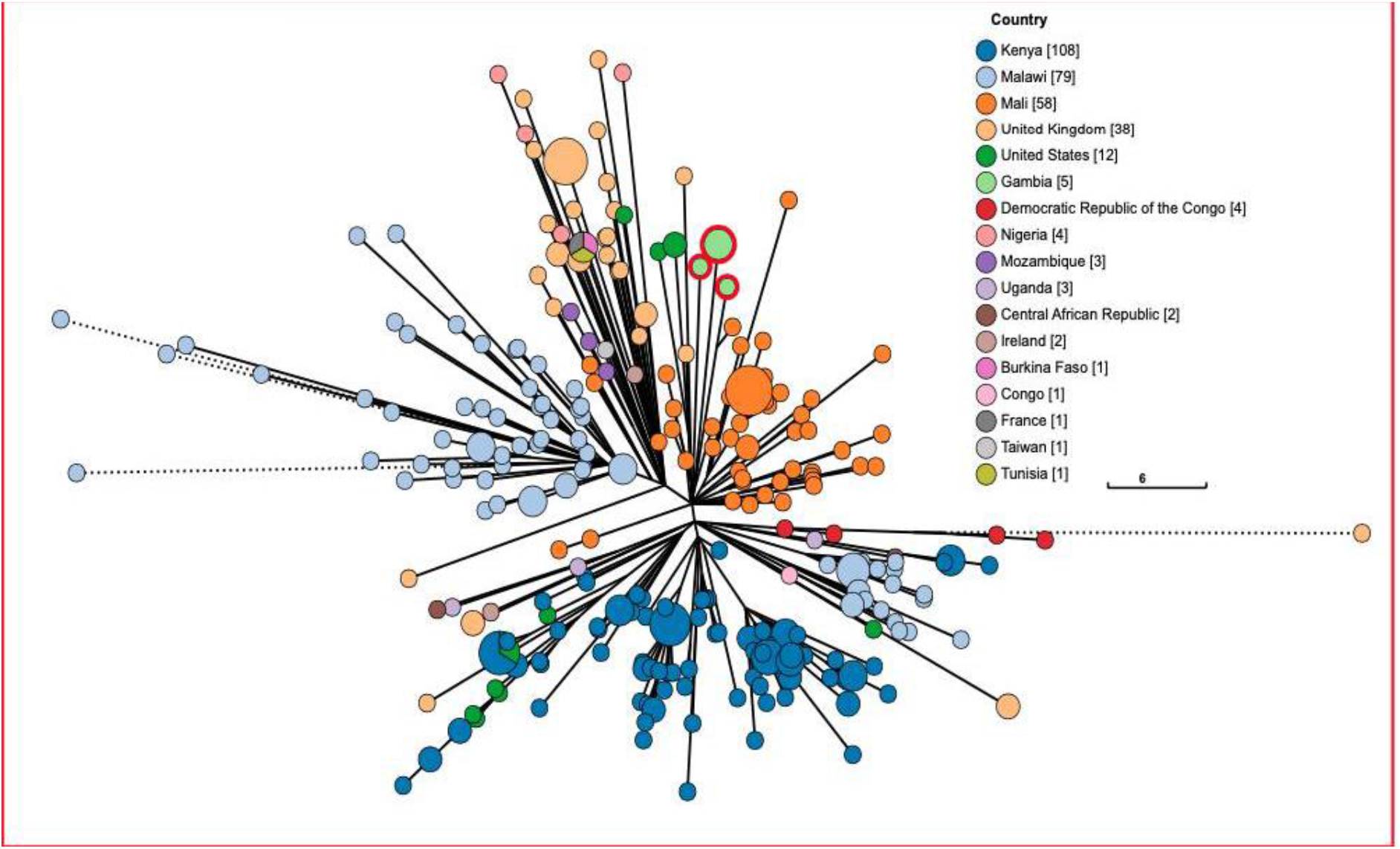
Phylogenetic tree of five *Salmonella* Typhimurium ST313 isolates from our study and all ST313 isolates from other countries (as indicated in the legend). Isolates from our study are highlighted in green with a red ring and are clustered away from the Kenyan (dark blue) and Malawian (sky blue) ST313 strains.

### Distribution of virulence, resistance and plasmid genes

A total of 124 virulence genes within and outside the *Salmonella* pathogenicity islands (SPI) were detected. The distribution of virulence genes detected and how they grouped based on the loci present can be found in Supplementary Table 1. Some virulence genes were conserved in the *Samonella* isolates evaluated while others were only present in some serovars. For example, SPI-7 which encodes *vex* and *tvi* genes was found in Typhi serovars only while SPI-11, which encodes the *CdtB* gene was found in several serovars within the atypical group.

Some genes found outside the SPI, including fimbriae and adhesion encoding genes as well as the type 1 fimbriae, were conserved in all isolates. Most of the genes that were variable in their distribution were found residing outside the pathogenicity islands. These genes included Gifsy-1 found in Typhimurium and Paratyphi C serovars only, and Gifsy-2 effector genes found only in Bron, Dublin, Enteritidis, Paratyphi C and Typhimurium isolates. Interestingly, we found 42% (31/68) of serovars in the atypical group had the virulence gene *cdtB* and that this gene was present in all our Typhi isolates.

Genomic analysis indicated more antimicrobial resistance genes in Enteritidis than any other serovar. Analysis of phenotypic data showed a similar pattern where 80% to 100% (n=40) of Enteriditis isolates were resistant to all the antimicrobials tested except ciprofloxacin. 100% (n=40) sensitivity was observed in all Enteriditis isolates tested against ciprofloxacin (Figure 8A). Some of the resistance genes present in Enteritidis were also found in Typhimurium ST313 isolates, but were present in only few of the atypical serovars. All Typhimurim isolates (n=16) tested were resistant to gentamycin. We found only few plasmid genes in our dataset. This was more pronounced in some serovars such as Dublin, Enteritidis and Typhimurium. In fact, none of the Typhi strains had a plasmid gene and only a few of the atypical serovars had one or two plasmids. We found that some plasmids were specific to particular serovars. For example IncX1 was found only in Dublin isolates. IncFIIB was common in Typhimurium isolates while Incl1 and IncQ were found in all Enteritidis isolates (see Table 5 for full summary).

**Figure 8.**
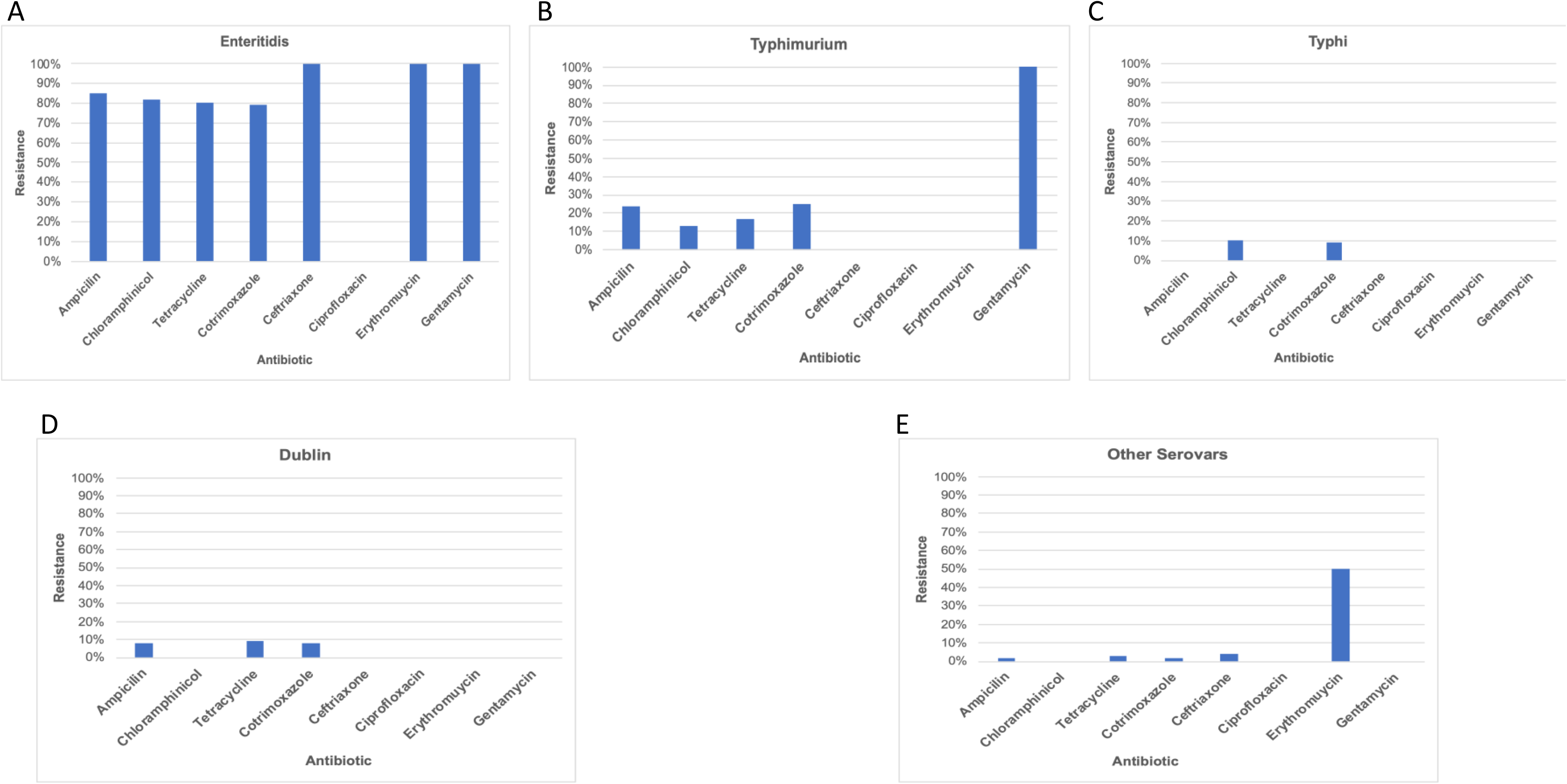
Antibiotic resistance patterns in invasive *Salmonella* serovars isolated in rural Gambia between 2008 and 2016.

## Discussion

In The Gambia, NTS is an important cause of invasive bacterial infections especially in children^14,34–37^. Using population-based epidemiological data and whole genome sequencing, we found an increase in the proportion of atypical NTS serovars causing invasive disease in rural Gambia between 2008 and 2016. We also observed changes in the incidence of disease over time. We identified sets of virulence genes in atypical serovar isolates that may be responsible for the increased prevalence of these serovars.

Few studies have described the distribution of non-typhoidal *Salmonella* serovars in The Gambia^14,37^. Between 2000 and 2004, Ikumapayi *et al*., reported Enteritidis as the major cause of invasive disease in rural Gambia while Typhimurium and other serovars accounted for only few cases^14^. Interestingly, the present study showed a significant reduction in the proportion of invasive *Salmonella* disease caused by Enteritidis. To identify serovars, Ikumapayi *et al*., used conventional antisera agglutination methods while polymerase chain reaction (PCR) methods were used for MLST typing^14^. This could underestimate the proportion of some serovars as antisera-based methods are limited in their ability to distinguish between closely-related and polyphyletic serovars^38^. By exploiting the advantages of whole genome sequencing, we identified 31 different serovars and thus a greater diversity of *Salmonella* serovars causing invasive disease. Between 2005-2015, Kwambana-Adams *et al*., reported Typhimurium to be the predominant invasive serovar in the coastal parts of The Gambia^37^, with 25% of isolates being serovars other than Typhi, Typhimurium, or Enteritidis. In comparison, our data show temporal and/or regional differences in the prevalence of *Salmonella* which could be attributed to many factors including host and pathogen genetic characteristics.

Globally, Typhimurium and Enteritidis are the two major serovars associated with invasive *Salmonella* disease^39,40^. However, this trend was different in rural Gambia where atypical serovars including Dublin, Virchow and Poona are increasing in prevalence. Studies have shown that genetic factors and immune status predispose individuals to invasive *Salmonella* disease^4^. For example, malnutrition and HIV have been associated with increased susceptibility to invasive *Salmonella* disease^41^. However, in The Gambia, the prevalence of malnutrition and HIV has not changed over the years suggesting that the increased incidence of invasive *Salmonella* disease may be attributable to other environmental factors or the genetic characteristics of the pathogen. We observed an increase in atypical serovars with the majority of cases occuring between 2012 and 2014. However, genomic analysis revealed various virulence factors implicated in invasion, proliferation and or translocation by Type III secretion systems in all Dublin isolates. Between 2012 and 2014, Dublin was the most common serovar isolated within the atypical group. Studies have reported that Dublin is associated with more severe disease and more frequently the cause of invasive disease than other types of non-Typhi *Salmonella*^42,43^. The present study reported two deaths associated with the Dublin serovar ranking second in mortality after Enteritidis. Moreover, this study identified the cytolethal distending toxin gene (*CdtB*) in the majority of atypical serovars (Clade G). This gene encodes cytolethal distending toxin (CDT) which activates host DNA damage and thus leads to G2/M phase arrest12. Analysis of all *Salmonella* genome assemblies in RefSeq (accessed 26-03-2020) showed overall prevalence of *cdtB* to be 35% (3832/10882), and when Typhi is excluded, this falls to 14% (1628/8678). This shows an uncommonly high level of *CdtB* in our atypical serovars. Experimental studies show that populations of HeLa cells infected with *cytolethal distending toxin* (CDT)-positive NTS serovars have a significantly larger proportion of cells with DNA damage response protein (53BP1) and γH2AX foci than CDT negative serotypes^12^. More importantly, *in vivo* analysis showed increased colonization of the host by CDT-producing pathogens that was associated with tumorigenesis and neoplastic lesions that led to chronic infections^12^. Thus, we speculate that increased prevalence of *cdtB* genes in our study may provide these serovars with a fitness advantage over Enteritidis and Typhimurium, potentially contributing to the shift we observed.

In contrast, we observed a high proportion of Enteritidis between 2010 and 2011. This period coincided with heavy rains resulted in severe flooding in the Upper River Region. Subsequent high rates of malaria infection may have influenced the population’s susceptibility to iNTS disease. Phylogenetic analysis of the Enteritidis isolates suggests a potential outbreak. All Enteritidis isolates recovered during this period were isolated within the Basse area with similar virulence and antimicrobial resistance patterns. Outbreaks of *Salmonella* Enteritidis as a result of consumption of contaminated food or animal products have been reported elsewhere^44^. Although this theory could be true, a study in Mali highlighted that, in contrast to *Salmonella* Typhimurium, iNTS disease caused by *Salmonella* Enteritidis started to increase from 2008 with the highest peak seen in 2010 and 2011^16^. The finding in Mali corresponds with our observed increase in Enteritidis in 2010 and 2011 suggesting the potential combination of a regional increase in Enteritidis exacerbated by the impact of the flood in our setting.

Antibiotic resistance in some *Salmonella* serotypes has been reported in many parts of Africa including The Gambia^14,45^. Our Enteritidis serovars had more resistance genes than other serovars. Similar findings were also reported in previous studies done in The Gambia which showed high percentages of multidrug resistance among *Salmonella* Enteritidis isolates^14^. However, five of our Typhimurium isolates of the ST313 genotype had resistance genes similar to those found in Enteritidis. In Kenya and Malawi, a distinct genotype of Typhimurium ST313 was reported to have a multidrug resistance gene located on a virulence plasmid^11^. Genomic analysis of all ST313 isolates in our study and those found in Enterobase suggest that this unique Typimurium ST313 is restricted to eastern Africa. Nonetheless, continued monitoring of these genotypes in other parts of Africa is vital. It is, however, reassuring that many of the atypical serovars did not acquire resistance genes, although continued monitoring is essential as antiomicrobial resistance (AMR) is increasing, and has a high global health burden. We found only one Dublin isolate with resistance genes.

## Conclusion

Overall, this study has shown a wide distribution of invasive *Salmonella* serovars circulating in The Gambia. More importantly, an increase over time in atypical serovars with high case fatality rates was also documented. The study highlighted the potential effect of some virulence genes in contributing to the shift we observed. However, experimental and functional studies could shed more light on the role of such virulence genes and the evolutionary pressures on these serovars. The shift in serovar prevalence could have implications for vaccine development and thus represent a public health concern. Therefore, investigations should be made to identify potential changes in the distribution of iNTS serovars elsewhere in Africa and the prevalence of these virulence elements.

## Authors and contributors

AK and GM conceived the research idea and AK wrote the first draft of the manuscript. AK, AP and NFA did the bioinformatics analysis. UNI, RS and JM did the microbiology. GD and team did the sequencing. AKS supervised AK and reviewed the manuscript. All authors have read and approved the final version of the manuscript.

## Ethical approval

The parent project consented participants before enrolling them in the study. Therefore, this study does not require any ethical approval.

## Data availability

The raw sequencing data is publicly available from the European Nucleotide Archive under BioProject PRJEB39996.

## Acknowledgements

We thank the following people: Gordon Dougan and team for sequencing the isolates. Thanh Le Viet for bioinformatics support, Abdul Khalid Muhammad and Nuredin Mohammed for statistics support and Jahangir Hussain for providing GEMS domestic data. AK also wishes to acknowledge the management of MRCG at LSHTM for partly funding his internship at Quadram while working on this project.

